# Behavior-Driven Marine Larval Dispersal and Settlement with AI Agent-Based Modeling

**DOI:** 10.64898/2026.04.29.721765

**Authors:** Xing Zhou, Guanghui Wang, Renzhi Wu, Annalisa Bracco

**Author notes:** Xing Zhou and Guanghui Wang contribute equally to this work. Renzhi Wu contributed in an advisory role only and was not involved in the experiments, data generation, or analysis.

## Abstract

Larval dispersal models are central to mapping and predicting ichthyoplankton dynamics in the ocean, yet despite decades of refinement they remain fundamentally limited by their ability to represent adaptive behaviors, relying instead on static trait parameterizations. This deficiency constrains our capacity to design effective restoration and mitigation strategies in an increasingly stressed ocean. SWARM (Simulating Waterborne Agent Routes for Marine connectivity) overcomes this barrier by integrating Large Language Model (LLM)-based behavioral agents with a standard biophysical model to simulate active decision-making during the pelagic larval stage. In both idealized and realistic conditions focusing on Red Snapper larvae in the Gulf of Mexico, agents develop adaptive behaviors that improve settlement and generate explainable vertical distribution patterns. SWARM demonstrates that LLMs can overcome long-standing limitations in dispersal modelling by explicitly representing behavioral drivers of movement, opening new pathways for predicting connectivity and designing effective marine-ecosystem restoration.

## Introduction

Ocean connectivity refers to the renewal, persistence, and recovery of marine populations by linking distant habitats through the movement of larvae, nutrients, and energy. Shaped by intertwined physical and biological processes, it is critical for larval dispersal, and a cornerstone of ocean ecosystem resilience, maintaining biodiversity and supporting the capacity of marine species to adapt to environmental stressors (Sponaugle & Cowen, 2009; Balbar and Metaxas 2019). For corals, fish, and many other taxa, this connectivity is largely set during the pelagic larval stage, usually indicated as pelagic larval duration (PLD), making larval dispersal a cornerstone of marine conservation and fisheries management (Selkoe & Toonen 2011; Treml et al., 2015; Torrado et al., 2021).

Biophysical models, typically implemented through Lagrangian particle-tracking, are foundational tools for studying marine dispersal (Lett et al., 2008; Paris et al., 2013). In these models, each particle represents an individual whose trajectory reflects both physical forcing (e.g., advection by currents) and organism-level behaviors. Growing evidence shows that larvae cannot be treated as passive tracers: their active behaviors strongly influence dispersal pathways, vertical positioning, and ultimately settlement success (Leis, 2020; James et al., 2023).

Conventional dispersal models encode larval behavior through static, empirically derived parameterizations (Swearer et al., 2019; Lombardo et al., 2022). For example, ontogenetic vertical migration (OVM) is often simulated using predefined probability matrices that cannot adjust to shifting environmental conditions (Karnauskas et al., 2022; Zhou et al., 2024; Bovee et al., 2025). These hard-coded rules, that for many traits are poorly known for most species (Moneghetti et al., 2019; Choukroun et al., 2025), overlook the flexible, context-dependent decisions larvae make in nature, limiting the realism and predictive power of traditional frameworks.

By contrast, replacing static parameterizations with adaptive, agent-driven behavior, can enable larval decision-making to emerge dynamically from environmental cues (e.g. temperature, light, and food availability) rather than being prescribed in advance. Large language model (LLM)-empowered AI agents excel in tasks that demand environmental sensing and adaptive decision-making, closely paralleling the complexities of larval behavior. For instance, embodied AI systems achieve robust navigation and manipulation in robotics (Yang et al., 2024; Mon-Williams et al., 2025), and AI-driven avatars in open-world games like Minecraft dynamically adjust their actions in response to changing surroundings (Smit et al., 2023; Wang et al., 2023). In the ocean context, an agent framework is particularly well suited for studying species with poorly understood biological traits and for exploring adaptive larval responses to environmental variability, as behavior emerges naturally from individual-level decision-making rather than being imposed a priori.

Here, we introduce SWARM (Simulating Waterborne Agent Routes for Marine Connectivity), a novel AI agent–based framework for modeling dispersal and settlement of marine larvae under time-varying, physically realistic ocean currents (Fig. 1, see also Methods). SWARM agents can infer behavioral responses from environmental cues, integrate text-based biological knowledge, and leverage memory of past environmental states and actions.

**Fig. 1.**
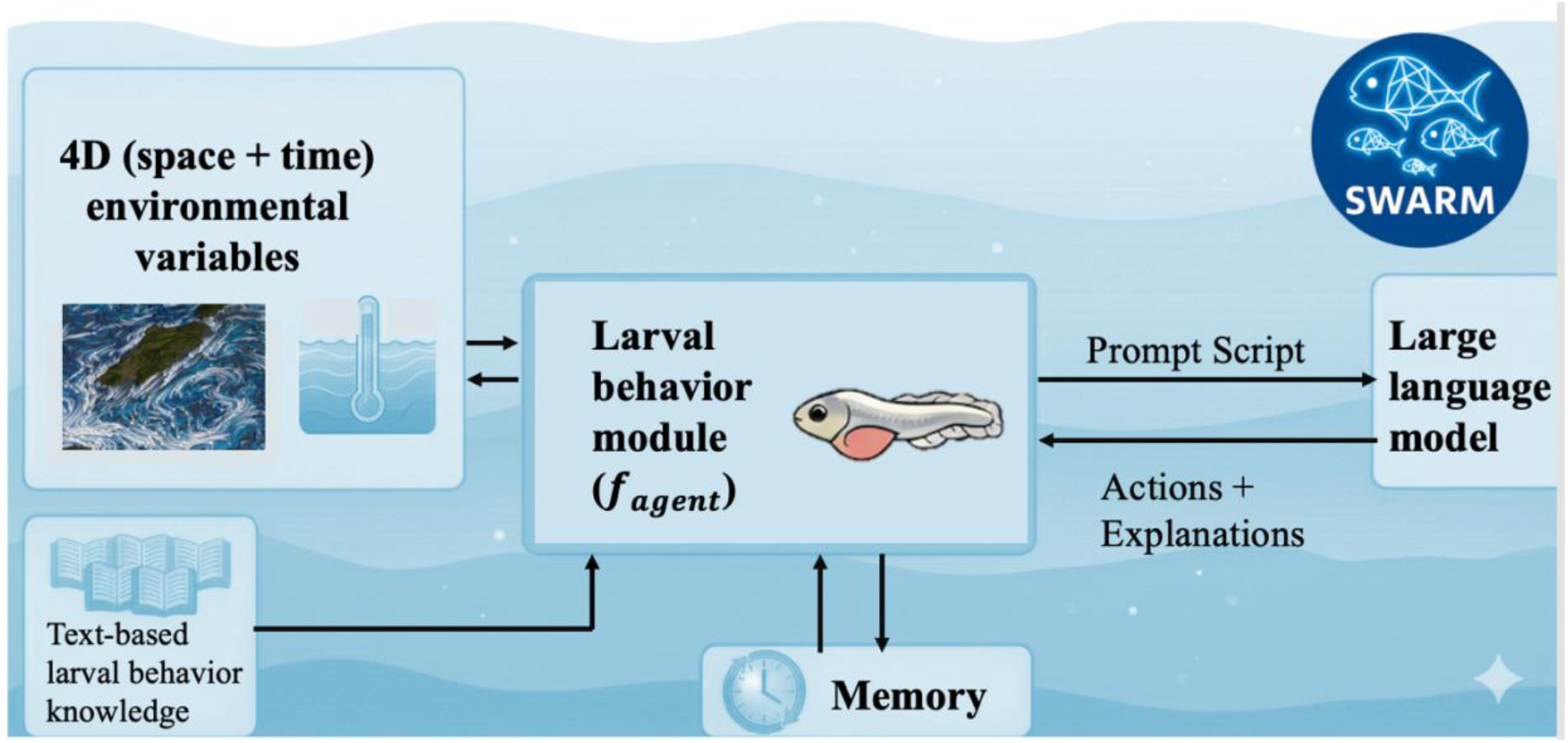
Schematic structure of SWARM. Initial template generated using Google Gemini (Google AI), with substantial modifications and annotations by the authors.

SWARM, coupled to a space-time representation of ocean currents and water characteristics (for example from an ocean model or a reanalysis dataset), provides a framework for investigating larval behavior in species with poorly constrained biological traits and for probing how larvae adapt to changing environmental conditions. We demonstrate its feasibility through an idealized experiment and a realistic regional application focused on Red Snapper larvae in the northern Gulf of Mexico (hereafter the Gulf). The Red Snapper is one of the most valuable commercial and recreational fish in the Gulf and SWARM provides novel insights into the role that OVM plays in its high ecological resilience (Erisman et al., 2020).

## Result

### Idealized Case

Figure 2A and 2B present the idealized set-up (see Methods) and the trajectories of all released larvae with the corresponding settlement rates for both mechanistic and SWARM approaches. Here, the settlement rate serves as a quantitative proxy for the relative importance of different behaviors in shaping larval connectivity.

**Fig. 2.**
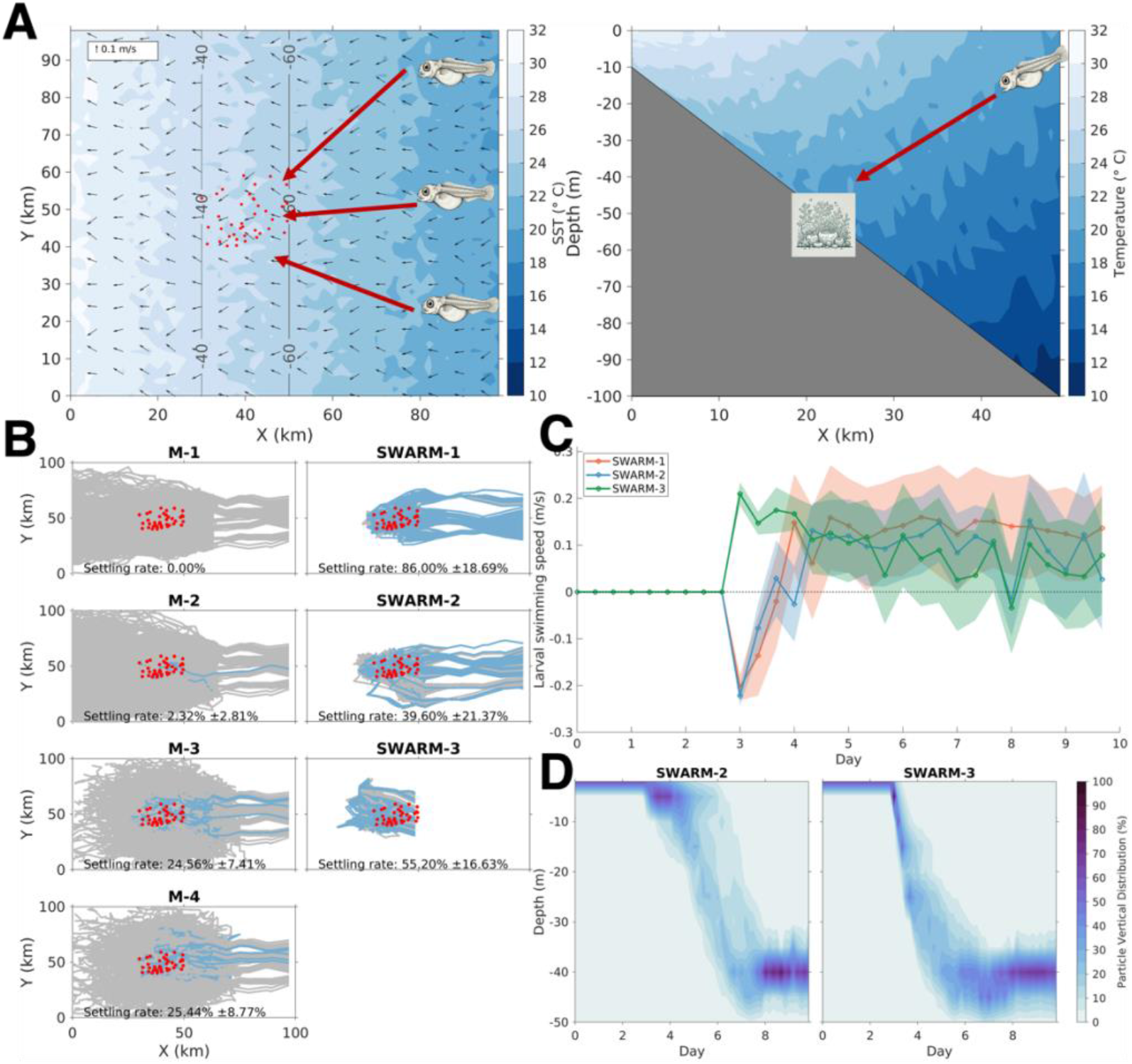
(A) Horizontal and vertical views of the virtual domain showing water temperature and surface currents. Red dots indicate coral reef locations, and gray contours denote the preferred settlement depth range of virtual larvae. (B) Trajectories of all released virtual fish larvae and corresponding settlement success rates (±1 SD) for experiments M1–M4 and SWARM-1–SWARM-3. (C) Larval swimming speed in the x (east–west) direction in SWARM agent-based simulations (positive values indicate eastward swimming against the current). Values are means across particles, with shading areas showing ±1 SD. (D) Emergent population-level vertical distributions from two AI agent-based simulations, resulting from individual larval decision-making.

The mechanistic experiments highlight the critical role of directional movement in larval settlement, supporting Berenshtein et al. (2022). In M-1, where no directional orientation was included, larvae were transported into nearshore areas, resulting in no settlement. Including reef-seeking orientation (M-2) increased the average settlement rate to nearly 2.3% with a subset of larvae that happened to pass through the coral reef zone resisting the currents and successfully settling. Adding rheotaxis orientation (M-3) and considering both rheotaxis and reef-seeking orientations (M-4), significant increase settlement rates to nearly 25%, and decreasing the length of the paths of the larvae. In essence, rheotactic orientation is the most influential behavior shaping larval dispersal and settlement outcomes in this idealized scenario.

Larval performance is substantially improved in all AI agent–based scenarios. SWARM trajectories closely resemble the successful one in M-3 and M-4 (short dispersal distances and direct movement toward the coral reef), with settlement rates significantly higher than in M-3 and M-4. The highest settlement rate is observed in SWARM-1, exceeding 86%. These results indicate that agent-based larvae exhibit adaptive decision-making, and their trajectories indicate strong west–east movement against currents.

The adaptive decision-making of the AI-based larvae is further illustrated by the swimming velocity in the x direction (Fig. 2C), which is positive throughout most of the time, indicating the emergence of rheotactic orientation across all simulations. Noticeably, though, the behavioral strategies diverge in each experiment. For example, in SWARM-3, the shift in spawning location causes the larvae to be transported to the western side of the reef region by the end of the egg stage, forcing them to swim aggressively eastward after days 3 and 4 (Fig. 2C, green line); and in SWARM-1 the swimming speeds are higher than in SWARM-2 during the settlement window, because in the latter, larvae can migrate into deeper waters where currents are weaker, thereby reducing the energy required to approach a suitable settlement habitat.

Another noteworthy outcome is the emergence of population-level vertical distribution patterns arising from individual larval decision-making (Fig. 2D). In SWARM-2, larvae remain near the surface for the first 6 days and then migrate to deeper depths in preparation for settlement; in contrast, in SWARM-3, larvae descend immediately after hatching and remain at approximately 40 m depth for most of their PLD. The difference in emergent vertical behavior is linked to the experimental configuration. The vertical distribution in SWARM-2 can be assimilated to a surface-dwelling strategy, commonly associated to species exhibiting long-distance dispersal, such as mojarras (Hernández et al., 2023), and is consistent with our experimental setup, in which larvae are spawned in the open ocean and later settle on shallower coral reef habitats. In contrast, the vertical distribution observed in SWARM-3 can be characterized as a deeper-dwelling strategy, commonly associated with coral reef fish species that show stronger local retention such as *Stegastes partitus* (Paris et al., 2004; Hernández et al., 2023).

### Red Snapper in the northern Gulf

Moving to the realistic Red Snapper case, Fig. 3A compares averaged vertical distribution in the Ichthyop and SWARM runs. SWARM larvae organize in vertical patterns similar to those imposed in Ichthyop.

**Fig. 3.**
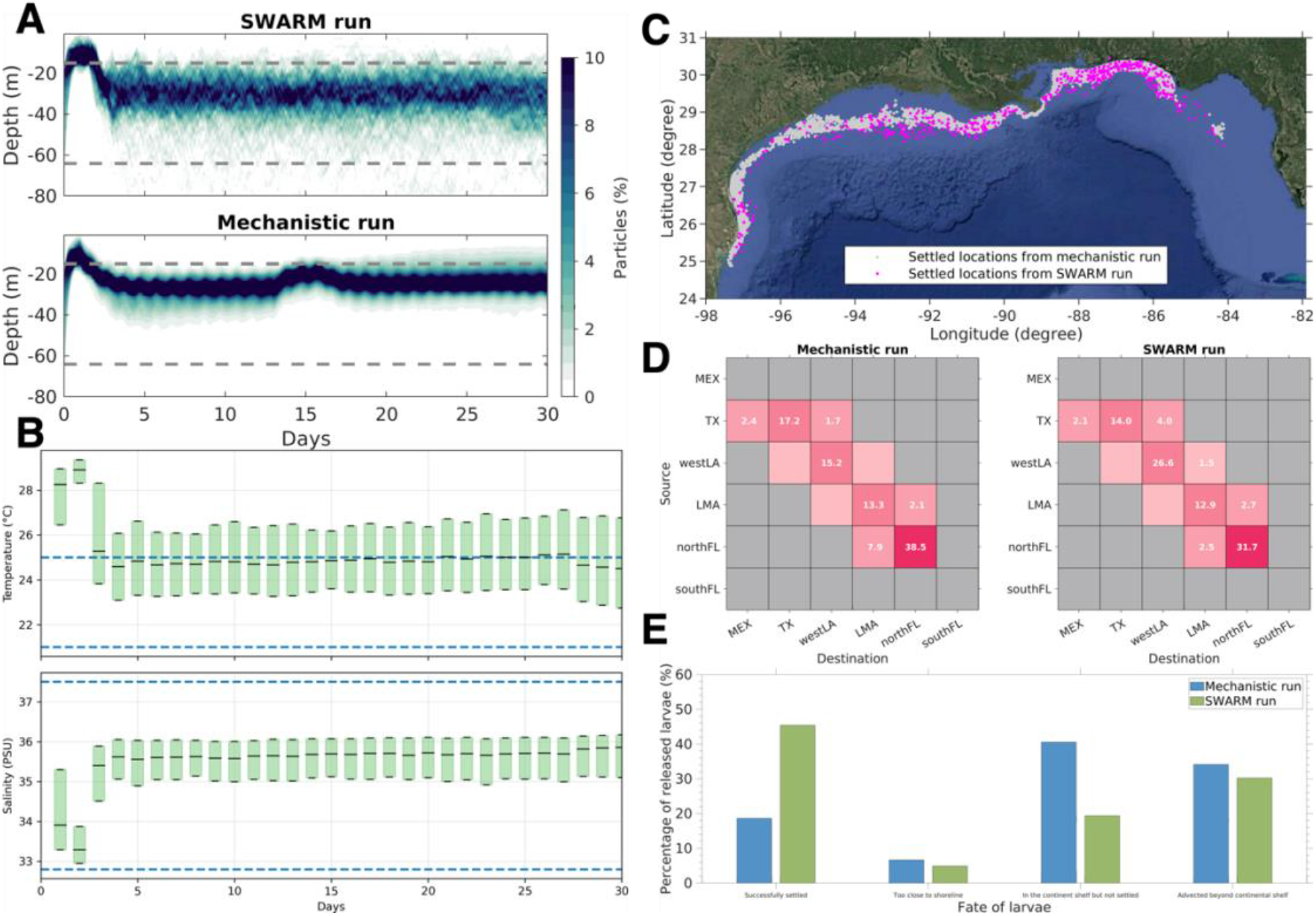
(A) Daily variation in the vertical distribution of Red Snapper larvae in the northern Gulf from the SWARM and mechanistic runs. Gray dotted lines indicate the 15–64 m depth range suitable for larval settlement. (B) Temperature and salinity conditions stored in agent memory in the SWARM run, summarized as daily whisker plots. Blue dotted lines indicate suitable ranges of temperature and salinity for Red Snapper larvae during the pelagic larval duration (21–25°C; 32.8–37.5). (C) Settlement locations of simulated Red Snapper larvae. (D) Connectivity matrices from the SWARM and mechanistic runs. (E) Bar plots illustrating the proportions of released larvae and their respective fates for the SWARM (green) and mechanistic (blue) simulations.

The agents inferred a downward movement after day 1 and maintained depths between 20 and 40 m during most of the PLD. A subtle divergence emerged only when approaching the settlement window, when agents tended to migrate deeper in preparation for settlement.

The analysis of the temperature and salinity states stored in the agent memory (Fig. 3B) indicated that temperature shaped this distribution. Agents moved deeper to avoid unfavorable surface temperatures during the first few days and subsequently remained near the upper temperature threshold, on the shallower side, to maximize food availability given that in our prompt design, water depth was used as a proxy for food resources.

Regarding settling locations and connectivity matrices, SWARM reproduced the patterns identified in the mechanistic runs (Fig. 3C and 3D). Both simulations showed strong local retention, with limited connectivity between regions west and east of the Mississippi Delta. Moreover, larvae spawned west of the Mississippi Delta preferentially settled closer to the coast than their release locations, whereas larvae spawned east of the Mississippi Delta showed a closer alignment between release and settlement locations.

The settling rate, however, diverge between the two runs, with SWARM showing more than a twofold increase in settlement (from 13% in the mechanistic run to ∼ 45%). Analyzing the fate of all released larvae (Fig. 3E), the proportions of larvae that were unsuccessful for being either too close to the shoreline or advected beyond the continental shelf (defined here as regions with water depth <150 m) were similar in the two simulations. This is because their positions are strongly modulated by physical advection. In SWARM, however, the fraction of larvae remaining on the continental shelf without settling is significantly reduced, as the adaptive behavior facilitates the “last mile” settlement process.

## Discussion

SWARM is a novel modeling framework that extends traditional parameterized representations of early life stages of marine organisms through the integration of LLM-augmented behavioral decision-making. A key advantage of SWARM is its ability to reduce the coding barrier associated with exploring and implementing larval behavioral traits in ecological models. In traditional approaches, customized behaviors need to be explicitly parameterized and hard-coded in rigid model structures. In contrast, SWARM allows behavioral logic to be specified through prompt scripts written in natural language, enabling users to incorporate complex behavioral hypotheses into numerical simulations in an accessible and transparent manner.

This accessibility is partly offset by current token costs, which limit the number of larvae that can be simulated. When agents interact with LLMs, token costs are incurred for both inputs and outputs. Input tokens include text-based larval behavioral knowledge, historical memory states, and current environmental conditions. Output tokens consist of the LLM’s internal reasoning, and the decisions returned to the model in JSON format. These costs increase with both the number of deployed agents and the communication between agents and the LLM. To ameliorate this issue, we implemented several strategies including batched API calls that integrate multiple agents into a single request and the use of structured system–user prompt designs. Despite these optimizations, the relatively high token costs of state-of-the-art LLM constrains the number of agents, and further reducing token costs remains a future objective. This could be achieved by developing smaller or task-specific LLMs, via fine-tuning or retrieval-augmented generation. Alternatively, domain-specific foundation models for marine organisms, trained on curated ecological and behavioral literature, could embed relevant knowledge directly in SWARM and improve efficiency. The rapid evolution of AI technologies will also ease this scalability concern.

A critical advantage of SWARM is that population-level patterns emerge naturally from individual decision-making. Instead of relying on fixed behavioral rules, SWARM allows larvae to respond autonomously to local environmental cues, producing realistic dispersal pathways, vertical distributions, and settlement patterns. This is an especially valuable feature for species with limited or uncertain behavioral data. Moreover, AI agents can be further augmented by explicitly constraining decisions through energy availability and metabolic expenditure, linking movement and habitat selection to physiological limits. This capability, combined with adaptive-making based on environmental cues and memory, offers a powerful avenue for exploring ecological responses to environmental stressors and climate change. As future conditions move outside the range of historical observations, empirical correlations become unreliable (Urban et al., 2016; Rose et al., 2024). By enabling agents to dynamically adjust behavior under novel environmental and energetic constraints, SWARM provides a flexible framework for probing organismal responses in a changing ocean and supports the development of restoration, management, and conservation strategies.

## Method

### Model Description

SWARM, written in Python and Java, links larval advection by ocean currents with a LLM accounting for their behaviors. In a typical Lagrangian-based biophysical modeling approach, the displacement of each virtual larva X can be described as:

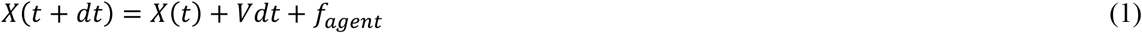

where *X*(*t* + *dt*) and *X*(*t*) represent the three-dimensional particle positions at times *t* and *t* + *dt*, respectively. *Vdt* accounts for displacements driven by hydrodynamic processes, such as advection by the ocean currents and diffusion. *f*_*agent*_, on the other hand, represents displacements due to larval behaviors. To ensure flexibility, *f*_*agent*_ has been coded as the sum of terms describing traits deterministic and/or well constrained for some species, such as buoyancy when well established, active swimming (*f*_*swim*_) and ontogenetic vertical migration (*f*_*OVM*_), and a module *f*_*agent*_, that enables interaction with a LLM (Fig. 1). *f*_*agent*_, receives user-defined environmental state variables (e.g., three-dimensional currents, temperature and salinity distributions in our next examples, but can also include nutrient concentrations, light or seabed type), together with a text-based prompt encoding behaviors to be considered. The prompt, tailored to each species, defines key responses to environmental conditions, habitat or settlement preferences, and may include active swimming and OVM if poorly constrained. These inputs are aggregated and communicated to the LLM via API (application programming interface) calls. In response, the LLM returns a JSON-formatted output containing the larval velocity components (*u*_*agent*_, *v*_*agent*_, *w*_*agent*_) in the *x, y*, and *z* directions due to their traits, along with the reasoning underlying the decision. The resulting action is then passed back to the Lagrangian model (Eq. 1) to advance the simulation, while the reasoning can optionally be stored for post analysis. This process is repeated for each virtual larva at user-defined time intervals. Another important component of *f*_*agent*_, is a memory module that stores historical environmental states and past decisions, enabling the LLM agent to learn over time and allowing it to refine future decisions, adapting dynamically to evolving environmental conditions.

### Implementation and Performance

The SWARM framework is publicly available on GitHub under an MIT license and is provided in two versions. The first is Python-based and is designed for rapid prototyping and testing of AI-agent behaviors. This version is intended for controlled experimentation with prompt design, decision rules, and larval–environment interactions and was applied in the idealized experiments presented in the next section.

The second version integrates SWARM directly into Ichthyop (Lett et al., 2008; Barrier et al., 2025) and is designed for real-world oceanographic applications. It supports high-performance simulations driven by realistic hydrodynamic fields with the possibility to enable or disable the AI-agent module through the Ichthyop graphical user interface. When activated, users provide an external JSON configuration file specifying the selected LLM and API credentials, together with a prompt script written in natural language to tailor agent behavior to the target organisms.

In terms of computational performance, a single iteration of the idealized case presented in this study requires approximately 1.5 hours when conducted on a modern laptop using a single CPU core. A 30-day regional application for the northern Gulf requires approximately 10 hours under similar conditions. The primary computational time arises from waiting for API calls, with an average estimated time of approximately 0.013 hours per call.

### Experiment design

We developed two test cases in an idealized and realistic ocean setting to evaluate whether the AI agents could autonomously infer behavioral strategies from environmental cues and their design is summarized below. The LLM of choice for SWARM is ChatGPT5, developed by OpenAI, which has demonstrated ability to understand and generate human-like text with accuracy, and reasoning across a wide range of tasks.

#### Idealized case

Some fish larvae use external cues for directional movement with the end goal of improving settlement rates (Leis, 2020; Dowine et al., 2021; Berenshtein et al., 2022). In an idealized environment where neglecting this trait leads to incorrect model predictions of dispersal and settlement, we compared SWARM and a traditional mechanistic model that explicitly included rheotactic orientation (the ability to swim against currents) and reef seeking orientation capabilities. We created a three-dimensional virtual domain measuring 100 km by 100 km horizontally, with a maximum depth of 100 m that featured strong east-to-west currents of approximately 0.1 m/s and random variability in the north–south direction (Fig. 2A). The domain extended from the nearshore at X = 0 km to the open sea (X = 100 km), with a temperature field at the surface that decreased linearly from 30°C nearshore to 20°C at the eastern boundary, modulated by random noise ranging between -1°C and 1°C to account for daily variations. The water depth increased uniformly from 10 m nearshore to 100 m in the open sea, and both temperature and current speed decreased with depth. Several coral reefs were placed at the center of the domain.

Virtual larvae were released at the open-ocean side uniformly between y = 30 km and y = 70 km. The PLD was set to 10 days, with the first three days spent in egg stage (larvae behaving as passive tracers), the following five days spent as larvae, and with a settlement window from days 8 to 10. We assumed an optimal temperature range of 22–27 °C for the larvae and a preferred settlement depth between 40 and 60 m. Settlement was considered successful whenever a larva could be found within 0.5 km of a coral reef.

A series of experiments was designed to validate SWARM (Table 1). In the mechanistic approach (M), we conducted four simulations, incorporating swimming behaviors with different orientation strategies. M-1 had no specific orientation (random swimming), while the other three incorporated reef-seeking orientation (M-2), rheotactic orientation (M-3), and a combination of both behaviors (M-4). The parameterized behaviors followed the implementation adopted in a widely used community model for simulating ichthyoplankton dynamics, known as Ichthyop (https://ichthyop.org/documentation/, accessed May 2025, Barrier et al., 2025).

**Table 1.** Description of experiments in the idealized case study.

| Experiments | Approaches | Behaviors |
| --- | --- | --- |
| M-1 | Mechanistic approach | Random swim |
| M-2 |  | Reef orientation |
| M-3 |  | Rheotaxis orientation |
| M-4 |  | Reef orientation + Rheotaxis orientation |
| SWARM-1 | 2D AI agent-based approach | Independently develop strategies |
| SWARM-2 | 3D AI agent-based approach | Independently develop strategies |
| SWARM-3 | Sensitivity test with spawning location shifted to X= 50 km (within the coral reef area). | Independently develop strategies |

We then conducted three AI agent–based simulations. In SWARM-1 vertical movement was not permitted, confining larvae to move at a fixed depth, to examine whether larvae developed more aggressive horizontal strategies. In SWARM-2 larvae were free to move in all three directions, to quantify how vertical migration impact emergent behavioral strategies, and lastly, in SWARM-3 the spawning area was changed to test the robustness of the agent emerging dispersal strategies.

The larval behavior knowledge provided to the agent included instructions for movement behaviors (vertical migration, active swimming, and settlement), as well as general guidance based on environmental cues (temperature, bathymetry, flow, and coral presence). For example, the agent was instructed to adjust its vertical position to remain within an optimal temperature range for survival, based on the general understanding that surface waters are typically too warm and very deep waters too cold.

Fish larvae often rely on chemical or acoustic cues to locate suitable settlement habitats (Montgomery et al., 2006; Gerlach et al., 2007) – in our case the reefs -, and we implemented a probabilistic cue field. We defined it using a spatially decaying probability function centered on the coral reefs, with a signal strength decreasing exponentially with distance, reaching 10% of its peak value at 2 km. The agent was instructed to respond to this gradient in its reef seeking behavior (see Supplementary Materials for full prompt script).

Since both mechanistic and SWARM approaches include a degree of randomness, we repeated the Ichthyop-based simulations 50 times, the AI agent–based simulations 10 times and analyzed the ensembles. The smaller ensemble size for SWARM was a trade-off between reducing token usage and achieving reliable simulation outcomes.

#### Realistic case: Red Snapper larvae in the northern Gulf

We further validated SWARM building upon the investigation of Red Snapper (*Lutjanus campechanus*) larval dispersal in the northern Gulf by Zhou et al. (2024). They adopted a mechanistic approach using the Ichthyop v3.3.17 model (Barrier et al., 2025) that incorporated egg buoyancy, OVM, and active swimming to explore the relative role of these traits and identified the OVM as a key trait shaping Red Snapper connectivity. In their simulations, egg buoyancy lifted larvae toward the surface during the hatching period, and OVM helped them toward their optimal settlement depths at later stages, substantially increasing settlement success. The OVM trait was imposed as a prescribed vertical distribution at different development ages following observations of the similar Mutton Snapper (*Lutjanus analis*) larvae (D’Alessandro et al. 2010) (direct observations of Red Snapper larval vertical distributions are lacking). In contrast, SWARM allows each larva to independently determine its vertical movement based on the local environment and historical memory, delivering an emergent population-level vertical distribution without prior assumptions.

For this work, we coupled SWARM to Ichthyop v3.3.17 and re-ran the connectivity scenarios under identical hydrodynamic forcing, replacing the behavioral parameterizations after egg stage with the LLM framework. Egg buoyancy was modeled deterministically, since it is governed by egg density and is a physical property. The hydrodynamic forcing was extracted from the realistic regional simulation of the Gulf circulation at 1 km horizontal resolution described in Zhou et al. (2024). We selected July 2016 among the two-year period (2015–2016) available because in this month large differences in settlement rates were observed between simulations that included or excluded OVM (Fig. S1a). We released ∼ 350 agents through a resampling procedure that preserved the initial release locations of the 21,000 virtual larvae in Zhou et al. (2024), as computational cost increases quasi-linearly with the number of larva-agents (Fig. S1b).

Red Snapper larvae were released according to the spatial distribution of adult Red Snapper (age > 2 years) following Dance and Booker (2019). Each larva was tracked for 30 days, with life history information provided through a prompt script available in the Supplementary Materials. Specifically, days 0–1 represented the egg stage, days 1–13 the pre-flexion stage (vertical movement only), days 13–26 the flexion and post-flexion stages (both swimming and vertical migration are possible), and days 26–30 the settlement window. All areas within the 15–64 m depth range in the northern Gulf were assumed to provide equal settlement opportunity. Successful settlement was defined as a larva being within 5 m of the seafloor within this depth range during the settlement window. In addition to life history information, the prompt included environmental guidance describing suitable temperature (21–25 °C, personal communication with Dr. Adela Roa-Varón) and salinity conditions (32.8–37.5 psu, Alan et al., 1980) for Red Snapper larvae, as well as water depth used as a proxy for food availability, to inform larval decision-making.

We compared the SWARM simulations with the most realistic configuration in Zhou et al. (2024), their Experiment 5, which included buoyancy, OVM, and swimming. In addition to vertical distributions, we assessed settlement locations and rates.

## Acknowledgements

This work was supported by a seed grant (GR00022420) co-funded by Microsoft and the Brook Byers Institute for Sustainable Systems and the Institute for Data Engineering and Science at Georgia Tech. G.W. was partially supported by the Apple Scholars in AI/ML PhD fellowship by Apple. We thank Dr. Jacob Abernethy for his mentorship and support.

## Author contributions

X.Z., G.W., R.W., and A.B. conceived the study. X.Z. and G.W. developed the SWARM framework and conducted the idealized case applications. X.Z. conducted the realistic regional application. X.Z., G.W., and A.B. contributed to writing, reviewing, and interpreting the manuscript. R.W. contributed in an advisory role. All authors read and approved the final manuscript.

## Code and data availability

The two versions of SWARM and the idealized test cases are available in the GitHub repository: https://github.com/SWARM-CODES/fish_llm

