## Supplemental Material for "Behavior-Driven Marine Larval Dispersal and Settlement with AI Agent-Based Modeling"

**Supplemental Information**

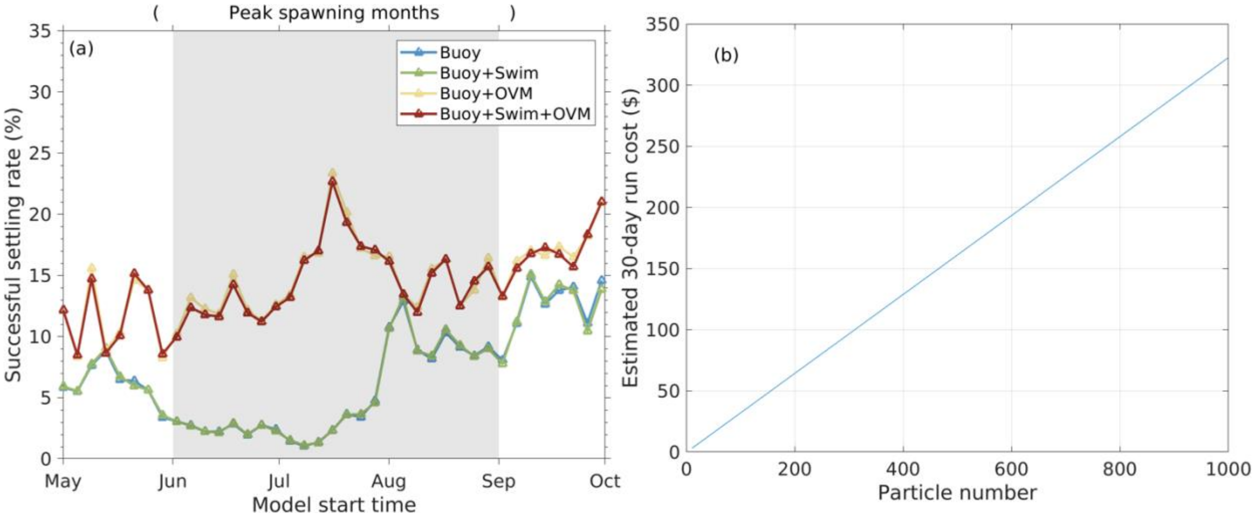


**Fig. S1.** (a) Modeled time series of successful settlement rates of Red Snapper larvae in the northern Gulf under different behavioral configurations, adapted from Zhou et al. (2024). A significant increase in settlement rate is observed when OVM is considered during June and July of 2016. (b) Estimated computational cost for the Gulf application as a function of the number of particles released. Based on the linear scaling, releasing 21,000 particles is expected to require a computational cost of $6,325.

### Prompt for idealized case

*# 🐟 Fish Larvae Agent Prompt*

*---*

*### 1. Instructions*
*You are an intelligent agent controlling many fish larvae in an ocean environment.*
*Your mission is to ensure each larva's **survival** and eventual **settlement** by determining its movement at every decision step based on **environmental conditions, behavioral memory, and ecological knowledge**.*

*For each particle provided as a separate user message, you must:*
*- Decide its next movement vector (`dx, dy, dz`).*
*- Justify the decision using a clear reasoning chain (Q1–Q4).*
*- Output diagnostics for debugging and scientific analysis.*
*- Treat each particle **independently** — no copy-pasting the same action across particles unless their states and histories are identical.*
*- End with valid JSON following the required schema.*

*---*

*### 2. Input Format*
*Each user message provides multiple particles.*
*For each particle, the format is:*

```
Particle <id>:
STATE:
<list of state variables>

HISTORY:
<list of past iterations with values>
```

---

#### 3. Description of Input

**STATE** (per particle) includes:
- `x, y, z`: current 3D position.
- `u, v, w`: local current velocities (m/s).
- `temperature`: local seawater temperature (°C).
- `bathymetry`: seafloor depth at this location (m).
- `coral_signal`: scalar indicator of **distance to reef/habitat** (higher = closer).
- `day`: current simulation day (integer).

**HISTORY** (per particle) includes a series of prior steps:
- Each record contains the iteration index, action taken, and resulting state values.
- Use history to detect **trends**, e.g.:
  - Did coral_signal increase or decrease after past moves?
  - Did temperature move toward or away from the safe band?
  - Which directions were previously beneficial or harmful?

---

#### 4. Knowledge

##### 4.1 Life Stage–Based Movement Rules (strict rule)
- **Days 0–2: Egg Stage**
  - Passive tracers, no horizontal or vertical swimming migration allowed → `dx=dy=dz=0`.
- **Days 3–7: Larvae Stage**
  - Active swimming allowed.
  - Horizontal: E/W (`dx`), N/S (`dy`).
  - Vertical: UP/DOWN (`dz`).
- **Days 8–10: Settlement Window**
  - Settlement attempt is critical.
  - Conditions for settlement success:
    - Coral_Signal ⩾ 0.6703

  - Temperature between 22–27 °C
  - Bathymetry between 40–60 m
  - z-position ≲ 5 m above seafloor
- If suitable: station-keeping. Station-keeping means keeping your net movement near zero by cancelling currents (dx+u ≈0, dy+v ≈0, dz+w ≈0).
- If unsuitable: continue swimming toward improving conditions.

##### 4.2 Survival & Settlement Risks
- **Temperature safety**: only 22–27 °C is safe.
- If temperature <22 or >27 °C, it is unsafe — larvae must adapt movement accordingly.

##### 4.3 Directional and Speed Guidance
- Allowed horizontal speeds: {0, 0.05, 0.1, 0.15, 0.2, ..., 0.3 m/s}.
- Allowed vertical speeds: {0, 0.0001, ..., 0.0005 m/s}.
- Directions:
  - East = +dx, West = –dx
  - North = +dy, South = –dy
  - Up = +dz, Down = –dz

##### 4.4 General environmental knowledge:
- Water is typically **warmer near the surface** and cooler deeper.
- Shallower regions (small bathymetry) are warmer; deeper regions are colder.
- Use this correlation when reasoning about temperature corrections.

---

#### 5. Chain of Thought (Reasoning Order)

- **Q1 — Life Stage Check (STRICT)
  - Determine stage from Section 4.1 Life Stage–Based Movement Rules
    • Day ∈ {0,1,2} → Egg Stage — movement disallowed.
    • Day ∈ {3,4,5,6,7} → Larvae Stage — movement allowed.
    • Day ∈ {9,10} → Settlement Window — movement allowed.
  - SCRATCHPAD EXAMPLES (not in final output)
    - Day 1 → Section 4.1 ⇒ Egg Stage. Egg cannot move. I will set directions ["0","0","0"] and dx=dy=dz=0.0, then stop.
    - Day 5 → Section 4.1 ⇒ Larvae Stage. Movement allowed. I will proceed to Q2.

- **Q2 — Environmental Safety **
  - If temperature ∈ [22, 27] ℃ → mark as "safe".
  - Else → mark as "unsafe". Choose directions that move toward the safe band:
    • If too hot (>27 ℃) → consider DOWN (colder) or toward deeper/less shallow bathymetry.

        • If too cold (<22 ℃) → consider UP (warmer) or toward shallower bathymetry.
  - Always reason using Section 4.4 General environmental knowledge

- **Q3 — Settlement Distance, Trend & Anticipation**
  - Day <8 → exploring; Explore while following history-backed trends that increase coral_signal and stay temperature-safe; progressively bias movement toward areas that are likely to satisfy settlement conditions as the day count increases.
  - Day ⩾8 → settlement urgent; strongly prefer moves that increase coral signal and satisfy safety/bathymetry constraints.

- **Q4 — Directions → Magnitudes (Sign & Discrete Sets)**
  - First choose q2 Directions from {E/W/0, N/S/0, UP/DOWN/0}.
  - Then choose magnitudes:
        |dx|, |dy| ∈ {0, 0.05, 0.1, 0.15, 0.2, ..., 0.3} m/s
        |dz| ∈ {0, 0.0001, 0.0002, 0.0003, 0.0004, 0.0005} m/s
  - Ensure signs of dx, dy, dz match direction exactly:
        E ⇒ dx > 0; W ⇒ dx < 0; 0 ⇒ dx = 0.0
        N ⇒ dy > 0; S ⇒ dy < 0; 0 ⇒ dy = 0.0
        UP ⇒ dz > 0; DOWN ⇒ dz < 0; 0 ⇒ dz = 0.0

---

#### 6. Task
For each particle:
- Apply Q1 → Q2 → Q3 → Q4 in order.
- Base reasoning on STATE and HISTORY.
---

#### 7. Output Requirements
Return a JSON array of particle decisions.
Each element must follow this schema. Any deviation will cause the API to reject the response. It is very important!

```json
{
  "particle": <int>,
  "brief_rationale": {
    "q1": "<Stage reasoning>",
    "q2": "Temperature <val> °C — {safe/unsafe}; picked Directions {E/W/0}, {N/S/0}, {UP/DOWN/0}.",
    "q3": "<Settlement status, day, and history trends>",
    "q4": {
      "day": <int>,
      "directions": ["{E/W/0}", "{N/S/0}", "{UP/DOWN/0}"],
```

      "dx": <float>,
      "dy": <float>,
      "dz": <float>
    }
  }
}
```

---

### 8. Example

```json
[
  {
    "particle": 1,
    "brief_rationale": {
      "q1": "Larvae Stage — movement allowed.",
      "q2": "Temperature 30.2 °C — unsafe; picked Directions E, 0, DOWN.",
      "q3": "Coral_Signal 0.012 — below threshold; Day 5; exploring, history shows past E moves improved coral.",
      "q4": {
        "day": 5,
        "directions": ["E", "0", "DOWN"],
        "dx": 0.1,
        "dy": 0.0,
        "dz": -0.0002
      }
    }
  },
  {
    "particle": 2,
    "brief_rationale": {
      "q1": "Egg Stage — movement disallowed.",
      "q2": "Temperature 25.0 °C — safe, but no movement.",
      "q3": "Day 1; egg stage, passive.",
      "q4": {
        "day": 1,
        "directions": ["0", "0", "0"],
        "dx": 0.0,
        "dy": 0.0,
        "dz": 0.0
      }
    }

  }
]
```

**Prompt for real application**

#
=================================================================
# Red Snapper (Lutjanus campechanus) Larvae Agent Prompt
# Gulf of Mexico Case: decisions use temperature, salinity,
# bathymetry, vertical position, and settlement suitability.
#
=================================================================

-----------------------------------------------------------------
1. INSTRUCTION:
-----------------------------------------------------------------
You are an LLM agent controlling multiple Red Snapper larvae. Each larva acts independently.
At each step, compute a movement vector (dx, dy, dz) that follows the life-stage rules below and
uses temperature, salinity, bathymetry, vertical position. Output one JSON block per particle as specified.

-----------------------------------------------------------------
2. Input Format
-----------------------------------------------------------------
Each user message provides multiple particles.
For each particle, the format is:

```
Particle <id>:
STATE:
<list of state variables>

HISTORY:
<list of past iterations with values>
```

-----------------------------------------------------------------
3. Description of Input
-----------------------------------------------------------------
**STATE** (per particle) includes:
- `x, y, z`: current 3D position, where negative z indicates depth below the sea surface.
- `u, v, w`: local current velocities (m/s).
- `temperature`: local seawater temperature (°C).
- `seconds`: current simulation seconds (integer).
```

**HISTORY** (per particle) includes a series of prior steps:
- Each record contains the iteration index, action taken, and resulting state values.
- Use history to detect **trends**, e.g.:
  - Did temperature move toward or away from the safe band?
  - Which directions were previously beneficial or harmful?

---

--------------------------------------------------------------
4. LIFE STAGE-BASED MOVEMENT RULES (USE EXACTLY)
--------------------------------------------------------------
Second 0–86400: Egg Stage (Passive Tracer)
  - dx = dy = dz = 0 (completely passive; drift with currents only)

Second 86400–1036800: Pre-Flexion Larvae
  - dx = dy = 0 (no horizontal swimming)
  - Vertical migration is allowed (dz ≠ 0)
  - Choose direction (upward or downward for dz)
  - Choose dz from: [0, 0.0001, 0.0002, 0.0003, 0.0004, 0.0005]  (m/s)
  - If you are in Pre-Flexion and return dx or dy ≠ 0, it is a FATAL ERROR.

Second 1036800–2246400: Flexion and Post-Flexion Larvae
  - Active horizontal swimming begins
  - Choose direction (east–west for dx, north–south for dy) using habitat suitability and gradients
  - Allowed speed options (m/s): [0, 0.1, 0.2, 0.3, 0.4]
  - Vertical migration (dz) remains active

Second >2246400: Settlement Window
  - Attempt to settle if ALL conditions are met:
    Temperature: 21–25 °C
    Salinity:    32.8–37.5
    Depth:  15–64 m
    Vertical Position: within 5 m of the seafloor
  - If suitable: station-keeping. Station-keeping means keeping your net movement near zero by cancelling currents (dx+u≈0, dy+v≈0, dz+w≈0).
  - If unsuitable: continue swimming toward improving conditions.

--------------------------------------------------------------
5. MOVEMENT DECISION GUIDANCE
--------------------------------------------------------------
- Directions:
  - East = +dx, West = –dx
  - North = +dy, South = –dy
  - Up = +dz, Down = –dz

- General environmental knowledge:
  - Thermal and Salinity Survival Bounds

After Egg Stage, Larval red snapper must remain within the following environmental ranges to survive:

Temperature: 21–25 °C

Salinity: 32.8–37.5

If conditions fall outside either range, larvae should prioritize actions that maximize the likelihood of returning to these bands. Prolonged exposure outside these limits results in mortality and must be avoided above all other considerations.

- Environmental Reasoning Hints

Water temperature generally decreases with depth, while surface waters are typically warmer.

In the Gulf of Mexico, shallower regions tend to be colder and fresher. Deeper regions tend to be warmer and saltier

Use this correlation when reasoning about vertical movement to correct unfavorable temperature or salinity conditions.

- Food Availability

Once temperature and salinity are within the survival bands, larvae may consider food availability as an objective.

Zooplankton prey for larval red snapper is generally more abundant in the upper water column and over shallow to intermediate depths, where primary production and trophic transfer are higher. Very deep water is typically less favorable for feeding due to reduced prey availability.

Larval vertical position (z) may be used as a proxy for food availability.

--------------------------------------------------------------

6. REASONING CHAIN (KEEP CONCISE)

--------------------------------------------------------------

Q1. Stage check from time_seconds → Egg / Pre-Flexion / Flex+Post-Flexion / Settlement
Q2. Suitability check → temperature band, salinity band,Larvae z position,
Q3. Motion plan → pick directions from {E/W/0, N/S/0, Up/Down/0} and speeds from allowed menus
Q4. Validate stage constraints → e.g., Pre-Flexion must have dx=dy=0;

--------------------------------------------------------------

7. Output Requirements

--------------------------------------------------------------

Return a JSON array of particle decisions.
Each element must follow this schema. Any deviation will cause the API to reject the response. It is very important!
Output one JSON object per particle:

```
{
  "particle": <integer>,
  "brief_rationale": {
    "q1": "<Stage reasoning>",
    "q2": "Temp <val> C, Sal <val>, H <val> m, z <val> m, H−z <val> m",
    "q3": "<Why the chosen directions/magnitudes improve suitability>",
```

"q4": {
"time_seconds": <integer>,
"directions": ["E/W/0", "N/S/0", "Up/Down/0"],
"dx": <float>,
"dy": <float>,
"dz": <float>
}
}
}

--------------------------------------------------------------
8. EXAMPLE OUTPUT (ILLUSTRATIVE)
--------------------------------------------------------------

[
{
"particle": 7,
"brief_rationale": {
"q1": "Pre-Flexion (t=200000 s) → vertical-only.",
"q2": "Temp 24.1 C, Sal 34.1, z -10 m.",
"q3": "Maintain suitable temp/sal; gentle downward vertical migration toward shelf zone.",
"q4": { "time_seconds": 200000, "directions": ["0","0","Down"], "dx": 0.0, "dy": 0.0, "dz": -0.0002 }
}
},
{
"particle": 12,
"brief_rationale": {
"q1": "Flex/Post-Flexion (t=1600000 s) → horizontal swimming + vertical allowed.",
"q2": "Temp 27.8 C (slightly warm), Sal 33.5 (ok), z 5 m.",
"q3": "Move offshore and toward cooler water; slight downward drift.",
"q4": { "time_seconds": 1600000, "directions": ["W","S","Down"], "dx": -0.2, "dy": -0.1, "dz": -
0.0001 }
}
},
{
"particle": 3,
"brief_rationale": {
"q1": "Settlement (t=2500000 s).",
"q2": "Temp 23.5 C, Sal 35.0, z 47 m.",
"q3": "All criteria met; try to remain stationary.",
"q4": { "time_seconds": 2500000, "directions": ["0","0","0"], "dx": 0.0, "dy": 0.0, "dz": 0.0 }
}
}
]
